# Zebrafish *acta1b* as a Candidate for Modeling Human Actin Cardiomyopathies

**DOI:** 10.1101/2023.08.29.555390

**Authors:** Kendal Prill, Matiyo Ojehomon, Love Sandhu, Sarah Young, John F. Dawson

**Author notes:** These authors contributed equally to this work (Kendal Prill, Matiyo Ojehomon).

## Abstract

Heart failure is the leading cause of mortality worldwide, primarily associated with cardiovascular disease. Many heart muscle diseases are caused by mutations in genes that encode contractile proteins, including cardiac actin mutations. Zebrafish are an advantageous system for modelling cardiac diseases due to their ability to develop without a functional heart throughout embryonic development. However, genome duplication in the teleost lineage poses a unique challenge by increasing the number of genes involved in heart development. Four actin genes are expressed in the zebrafish heart: *acta1b, actc2*, and duplicates of *actc1a* on chromosomes 19 and 20. In this study, we characterize the actin genes involved in early zebrafish heart development using *in situ* hybridization and CRISPR targeting to determine the most suitable gene for modelling actin changes observed in human patients with heart disease. The *actc1a* and *acta1b* genes are predominantly expressed during embryonic heart development, resulting in severe cardiac phenotypes when targeted with CRISPR. Considering the duplication of the *actc1a* gene, we recommend *acta1b* as the best gene for targeted cardiac actin research.

## Introduction

Heart failure is the leading cause of death worldwide, with cardiovascular disease being a major contributor [1]. Cardiomyopathies are referred to as “diseases of the sarcomere” because mutations in genes encoding proteins of the sarcomere contractile machinery are a primary cause of cardiomyopathies. These genes include myosin, troponin, tropomyosin, cardiac myosin binding protein C, and cardiac actin (ACTC) [2,3]. Recent efforts have targeted some of these proteins for drug development to treat cardiomyopathies [4].

Testing drugs within a whole animal system is crucial for developing treatments for diseases. One of our aims is to understand how changes in the cardiac actin gene (ACTC) lead to different cardiomyopathies. We have conducted molecular-level studies on several ACTC variants [5–9], however, our ultimate goal is to integrate our molecular knowledge of ACTC biochemical changes with physiological dysfunction by editing the cardiac actin gene in an animal model.

The zebrafish serves as an excellent model for cardiac research [10,11]. Their embryos are transparent, allowing detection of heartbeats at 24 hours post-fertilization (hpf), and they do not require a fully functional heart for viability for the first 5 days post-fertilization (dpf) due to oxygen diffusion through their tissues. Therefore, zebrafish offer an accessible model system to study embryonic lethal heart mutations found in mammals. Additionally, zebrafish are small, easy to maintain, exhibit rapid growth, have large numbers of progeny, and the genome of the Tübingen (Tü) strain has been completely sequenced.

However, unlike humans, which possess a single α-cardiac actin gene (ACTC), the zebrafish genome contains four actin genes expressed in the heart (referred here to as zfactc genes): *acta1b, actc2* (formerly referred to as *actc1c*), and identical copies of *actc1a* on chromosomes 19 and 20. Previous studies have explored the cardiac-related effects of mutations in *actc1a* and *acta1b* [12–14]. Furthermore, we identified and conducted preliminary characterization of *actc2* [15](also known as *acte1* [16]).

Shih et al. (2015) performed transcriptome analysis of *actc1a* and *acta1b* in zebrafish hearts at embryonic (96 hpf) and adult (6-month-old) stages, demonstrating that both paralogues are expressed in the heart at each stage. However, *acta1b* exhibited predominant expression in embryonic hearts, while *actc1a* was the dominant paralogue expressed in adulthood, suggesting developmental regulation of zfactc genes [16].

While early expression of *actc1a* and *acta1b* has been characterized using *in situ* hybridization (ISH), previous work primarily focused on the somites of the tail. The expression characterization of *actc1a* included the somites and the heart from the 1-4 somites to 7 dpf stages [16–18], while *acta1b* ISH analysis focused on the somites [18–20]. In our preliminary characterization of *actc2*, we observed its expression in the heart and somites at 36 hpf [15]. Due to the teleost genome duplication, the *actc1a* genes located on chromosomes 19 and 20 share identical sequences making it extremely challenging to design *in situ* hybridization probes or CRISPR sgRNA that can distinguish between the duplicate genes.

To determine the optimal ortholog among the zfactc genes for editing and modelling human cardiac diseases resulting from ACTC mutations, we studied the spatiotemporal expression of these genes in the heart and observed the functional consequences of CRISPRs targeting these genes. All three zfactc genes are expressed in the heart during the initial stages of heart development. However, the *actc1a* and *acta1b* genes are the dominant paralogues expressed during embryonic heart development and result in characteristic cardiac phenotypes when modified with CRISPR/Cas9. The *actc2* gene appears to be a minor contributor, with its cardiac expression primarily occurring in the first 48 hpf, and no observable *actc2* expression in the heart at 72 hpf. Given the duplication of the *actc1a* gene and the challenges associated with ensuring mutations in both copies (*actc1a* 19 & *actc1a* 20), we suggest that the *acta1b* gene is the best candidate for cardiac actin research.

## Materials and Methods

### Ethics Statement

All protocols were carried out according the guidelines stipulated by the Canadian Council for Animal Care and the University of Guelph’s Office of Research Animal Care Committee (Animal Use Protocol license: 4309)

### Zebrafish Maintenance

Adult zebrafish (Tübingen and TL/AB strain) were maintained according to guidelines by the Canadian Council on Animal Care and kept on a 12/12-hour light and dark cycle at 28°C. Adults were fed brine shrimp (Hikari Bio-Pure Brine Shrimp) and fish flakes (Omega One) daily in a cycled-water aquatic facility. Embryos were collected from crossing wild-type adult zebrafish and grown at 28°C in zebrafish embryo medium [21] for up to 6 days prior to fixation.

### In situ hybridization

Zebrafish embryos were staged and fixed in 4% paraformaldehyde/PBS overnight at 4°C. Cardiac actin probes were synthesized (Bio Basic) and cloned into pUC57-Amp for probe synthesis (Table 1). Antisense RNA probes were synthesized from the pUC57 construct using SP6 RNA polymerase (Thermofisher). *In situ* hybridizations were carried out as previously described [22].

**Table 1.**
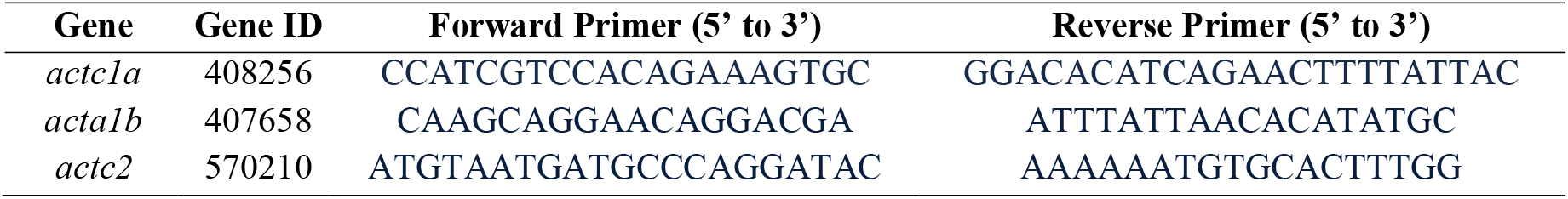
Primers used to amplify *in situ* hybridization probes.

### CRISPR sgRNA Preparation and Microinjections

CRISPR single guide RNA (sgRNA) was designed for *actc1a, acta1b* and *actc2* using CHOPCHOP (https://chopchop.cbu.uib.no; danRer11/GRCz11)[23], selecting the sgRNA that returned the fewest to zero off-target sites (Table 2). Given the extreme identity between the two *actc1a* genes physically located on chromosomes 19 and 20, sgRNAs were designed that targets both genes.

**Table 2:**
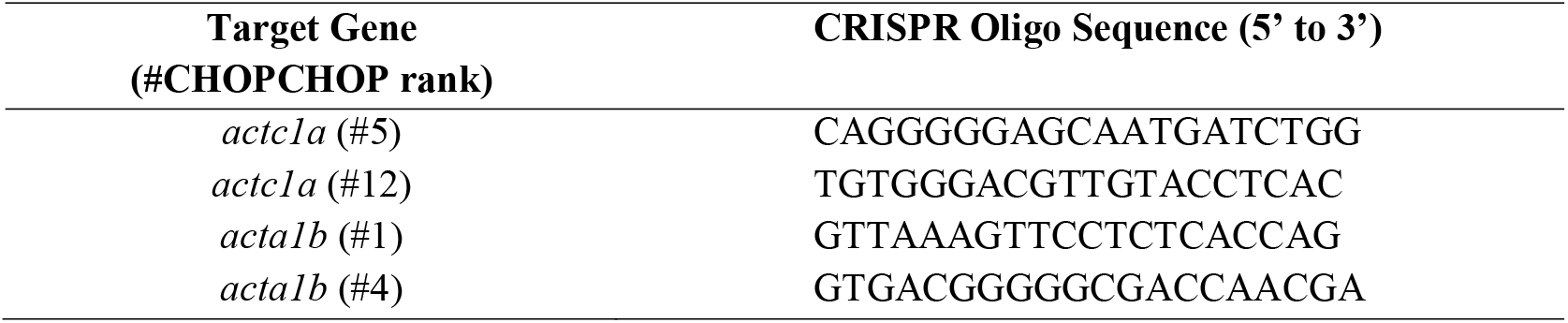
CRISPR single guide RNA designed to target the zfactc genes.

sgRNA was synthesized using SP6 RNA Polymerase (Thermofisher). Cas9 (1 mg/ml; CP01-200, PNA Bio Inc) and fresh sgRNA (1 ug) was injected into 1-cell zebrafish embryos that recovered in zebrafish embryo medium at 28°C. Zebrafish embryos were monitored daily for the appearance of phenotypes and imaged using an iPhone 6 camera (8 megapixel, 1080p HD video at 60 fps; Apple Inc.)

### High Resolution Melt Curve Analysis Screening

For screening using High Resolution Melt Curves (HRM), genomic DNA was extracted from zebrafish embryos by individually lysing tissue in 10 ul of 0.5 M NaOH at 95°C for 45 mins, followed by neutralization with 0.2 mM Tris-HCl (pH 8). DNA was diluted 1/20 for best amplification results during HRM. The HRM amplicons were designed to be no larger than 120 bp and centered on the CRISPR-Cas9 cut site (Table 3). High Resolution Melt Curves were produced using the saturating dye, EvaGreen (Type-it HRM PCR Kit, Qiagen) and the manufacturers recommended protocol for use. HRM was performed using a QuantStudio 7 Pro Real-Time PCR System (Thermofisher) and results interpreted using Design and Analysis Software 2.6 (Thermofisher) with the High Resolution Melt Analysis Module v2 (Thermofisher).

**Table 3:**
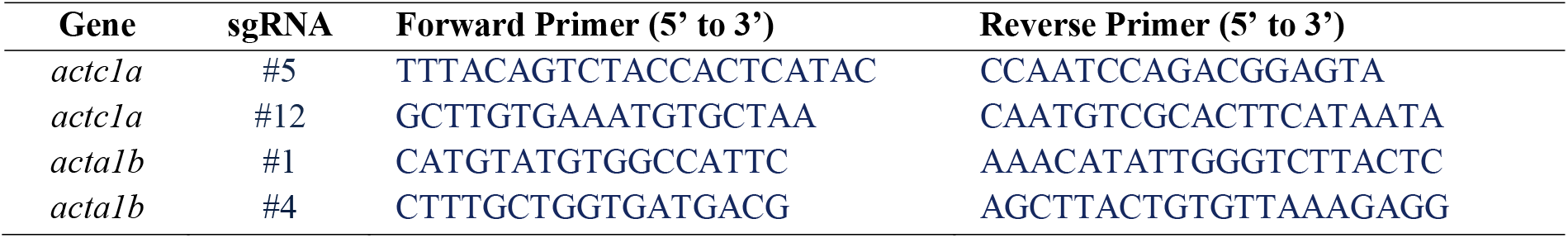
Primers employed for high resolution melting analysis of actin CRISPR amplicons.

### Heart Rate Acquisition and Analysis

The heart rates of all CRISPR-injected embryos were video-recorded daily from 1-5 dpf using an iPhone 6 camera (8 megapixel, 1080p HD video at 60 fps; Apple Inc.). Captured videos were analyzed with DanioScope software (Noldus) to determine heart rates. Genomic DNA was extracted from these embryos and screened for mutations using HRM. Embryos with mutated sequences had their heart rates graphed in comparison to wild-type controls.

## Results

### actc1a and acta1b are the predominant actin paralogues expressed during striated muscle development

To identify which actin paralogues are necessary for heart development, we analyzed the expression of *actc1a* (19&20), *acta1b*, and *actc2* using *in situ* hybridization (ISH) during the early stages of embryogenesis (Fig 1). At 24 hpf, all three cardiac actin paralogues were expressed in the linear heart tube (Fig 1, A-C), although *actc2* appears to be limited to ventricular cardiomyocytes (Fig 1C)[24]. *Actc1a, acta1b*, and *actc2* were all expressed in the embryonic zebrafish heart at 36 and 48 hpf, with the strongest expression observed in the ventricle (Fig 1). However, unlike *actc1a* and *acta1b, actc2* did not have any observable cardiac expression at 72 hpf, suggesting that *actc2* may only be required for the early stages of heart development and sarcomere assembly in ventricular cardiomyocytes.

**Fig 1.**
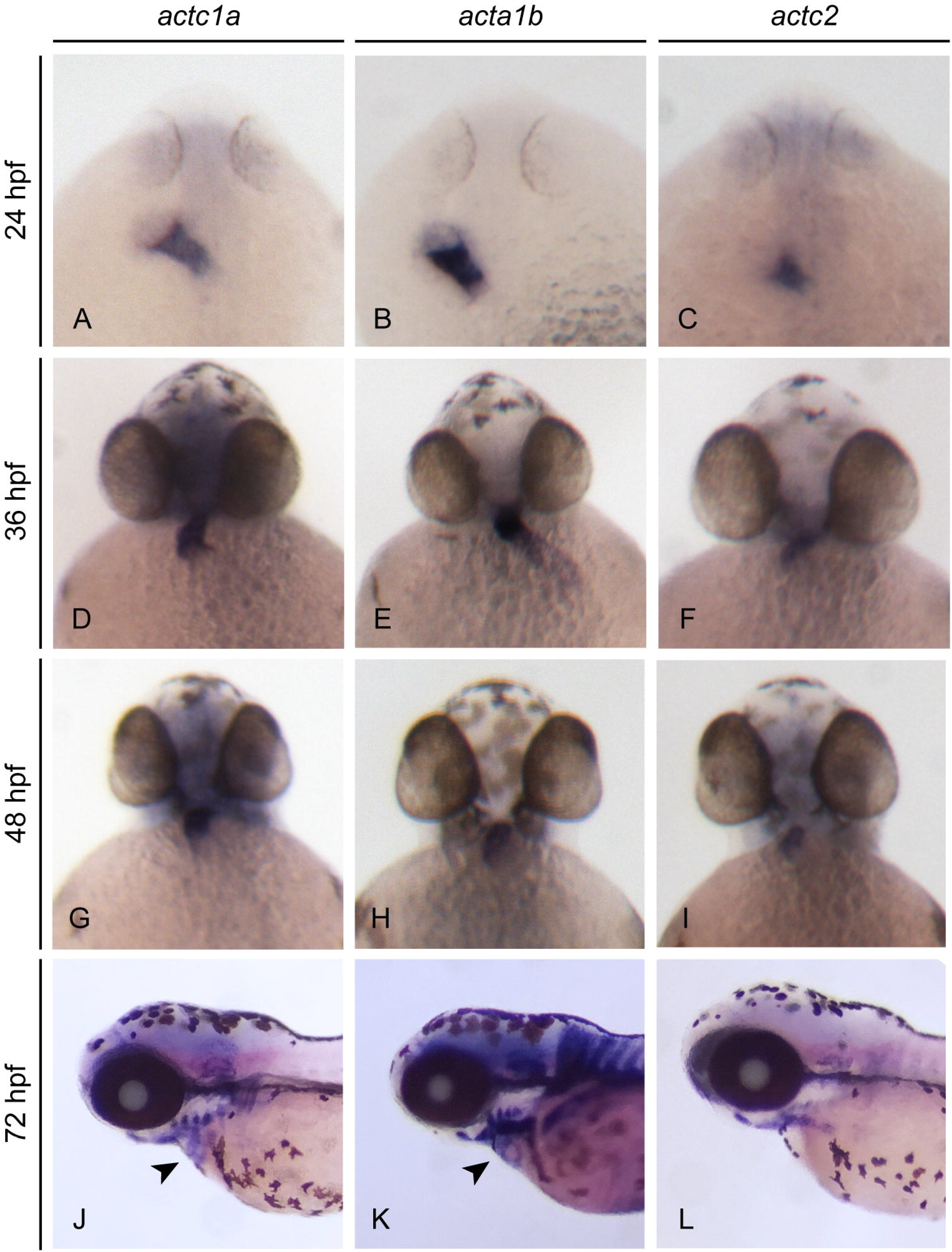
Expression of cardiac actin paralogues throughout early embryo heart development. At 24 hpf, *in situ* hybridization reveals heart tube-restricted expression of *actc1a* (A), *acta1b* (B) and *actc2* (C). By 36 hpf, *actc1a* (D) and *acta1b* (E) are expressed throughout the heart with the strongest expression observed in the ventricle. *Actc2* demonstrates an expression pattern that appears restricted to the outer curvature of the ventricle at 36 hpf (F). At 48 hpf, *actc1a* (G) and *acta1b* (H) are expressed only in the ventricle while *actc2* (I) maintains a small region of ventricle specific expression. *Actc1a* and *acta1b* have continued cardiac expression at 72 hpf (J&K, black arrowheads), while *actc2* (L) expression is no longer observed in the heart of developing embryos.

### actc1a and acta1b are required for normal heart development and function

We demonstrated that *actc1a, acta1b*, and *actc2* were expressed during the early stages of heart development at 24 hpf but exhibited different temporal and spatial expression patterns from 36-72 hpf (Fig 1). Since *actc1a* and *acta1b* demonstrated consistently strong expression in the developing embryonic zebrafish heart, we tested the necessity of these predominant actin paralogues in heart development by disrupting each one using the CRISPR/Cas9 system. We identified sgRNAs that specifically targeted *acta1b*, with no off-targets, and sgRNAs that targeted identical sequences in both copies of *actc1a* on chromosomes 19 and 20 (Fig 2A). We identified CRISPR-mediated cardiac actin mutants for analysis from wild-type (and uncut siblings) embryos by shifts in their sequence melting profiles (Fig 2B-E).

**Fig 2.**
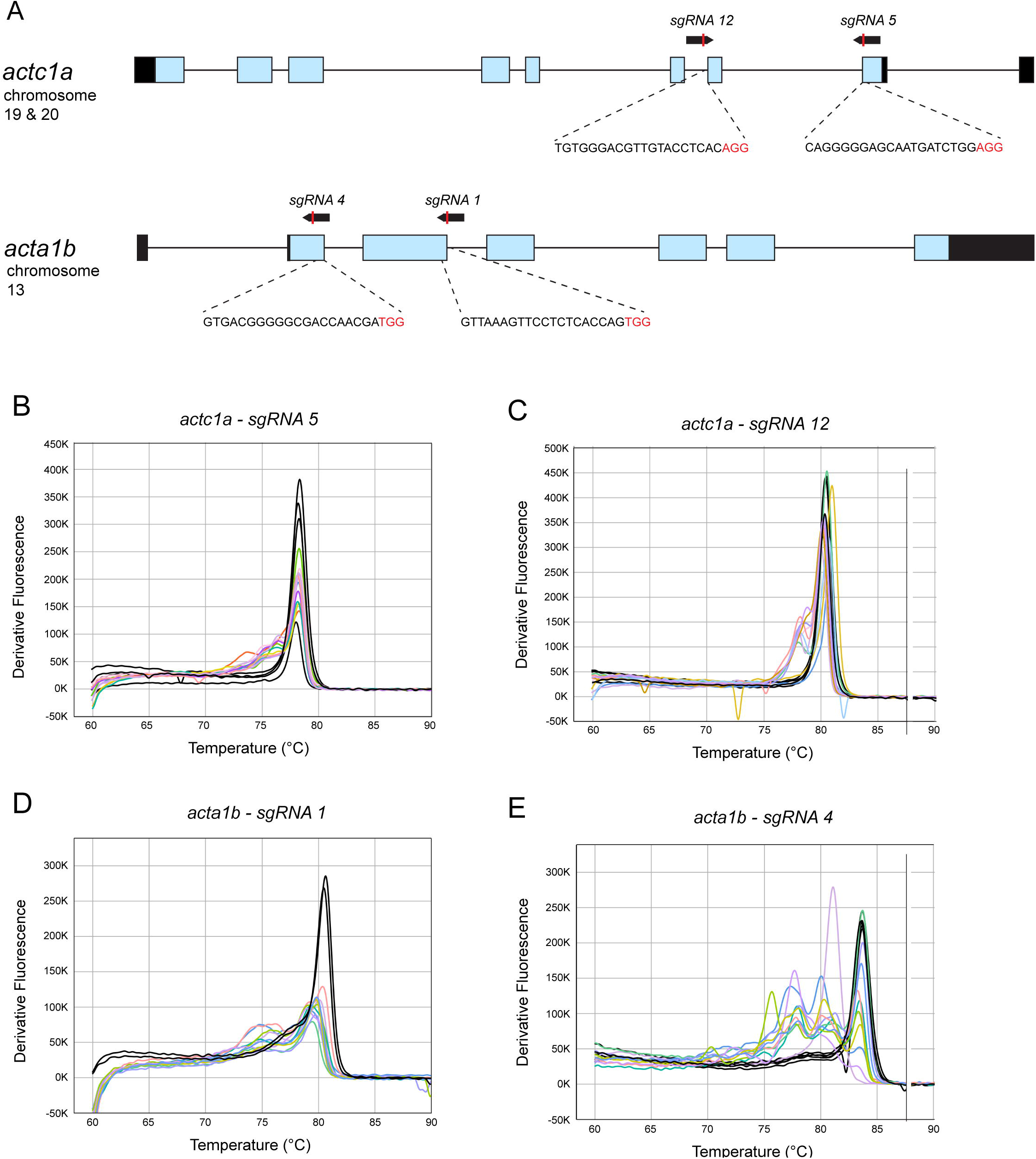
Actc1a and acta1b CRISPR target sites and HRM analysis. (A) A gene schematic view of *actc1a* (chromosomes 19 and 20), *acta1b* (on chromosome 13), and the CRISPR sgRNAs (shown below each gene) used to mutagenize each. CRISPR sgRNA sequences are shown in black, PAM sites in red. Melt Curves generated by HRM identified embryos with CRISPR-mediated mutations in *actc1a* (B&C) and *acta1b* (D&E) by differentiating the melting profiles of sequences with CRISPR-induced indels from wild-type. Black – wild-type; pastel – individual mosaic-CRISPR-mutants.

By 72 hpf, zebrafish embryos with *actc1a* or *acta1b* CRISPR-mediated mutations displayed cardiac phenotypes [25], including pericardial edema, unlooped hearts, enlarged chamber(s), degenerated hearts, and abnormal heart rates (Fig 3 & 4). The majority of embryos (89%) with *acta1b* CRISPR-mediated mutations showed significantly reduced heart rates compared to wild-type embryos at 72 hpf (Fig 4). *Actc1a* CRISPR-mediated mutants also displayed abnormal heart rates but showed an almost even distribution of normal (41%), higher (30%), and lower (28%) heart rates compared to wild-type among the CRISPR-mediated mutants at 72 hpf (Fig 4). By 120 hpf, a subset of *actc1a*-CRISPR and *acta1b*-CRISPR mutants displayed higher heart rates than wild-type embryos, suggesting a common compensatory mechanism for mutations in the dominant cardiac actins.

**Fig 3.**
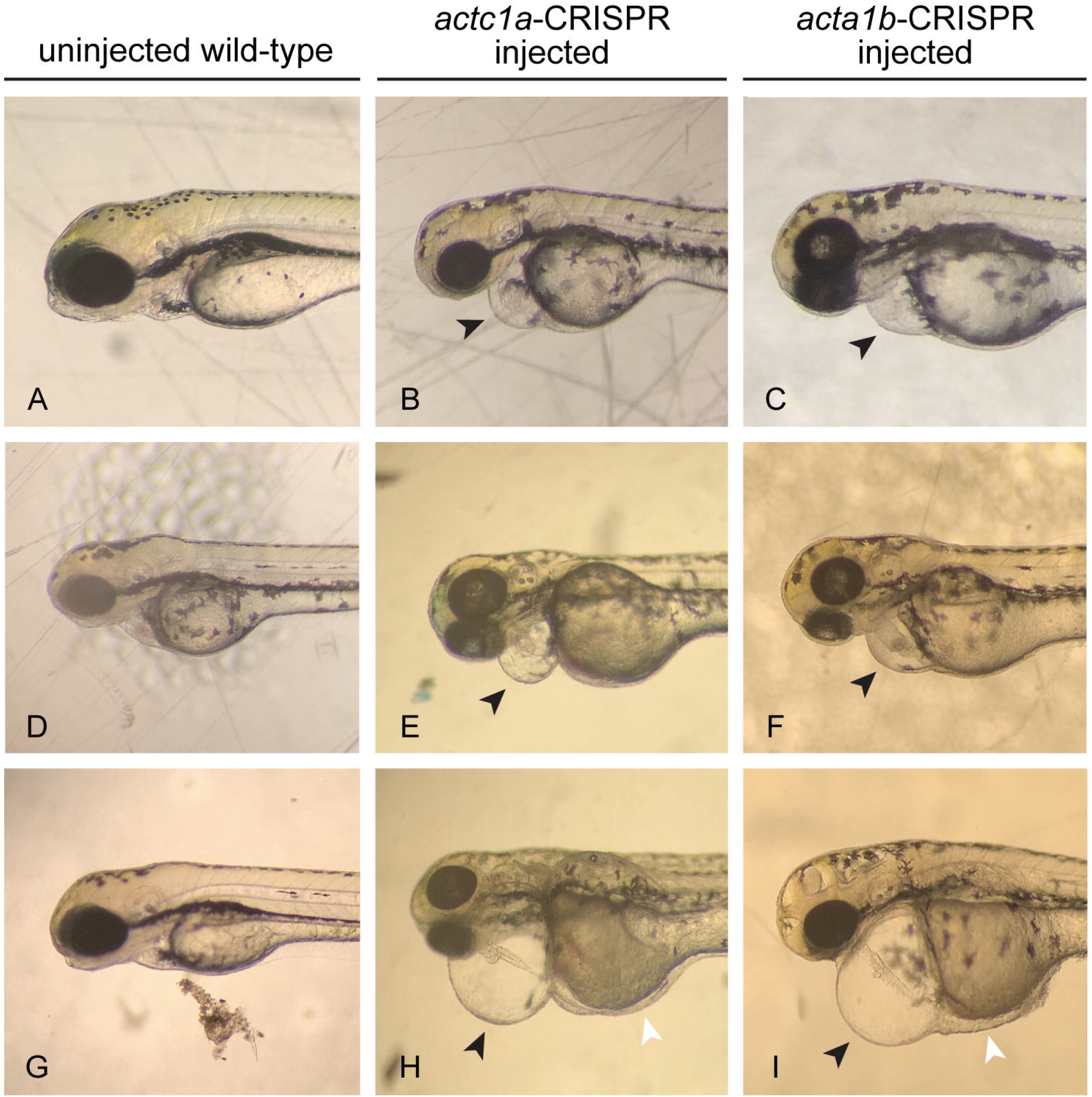
Phenotypes of cardiac actin CRISPR-injected embryos at 3 days post fertlization. When compared to wild-type control embryos (A,D,G), embryos injected with *actc1a* or *acta1b*-targeting CRISPR sgRNAs displayed pericardial edemas of varying severity (black arrowheads) and incompletely looped hearts (B,C,E,F,H,I). A subset of injected embryos, with CRISPR-mediated mutations, demonstrated extreme pericardial edema, yolk edema and degenerated hearts (H&I).

**Fig 4.**
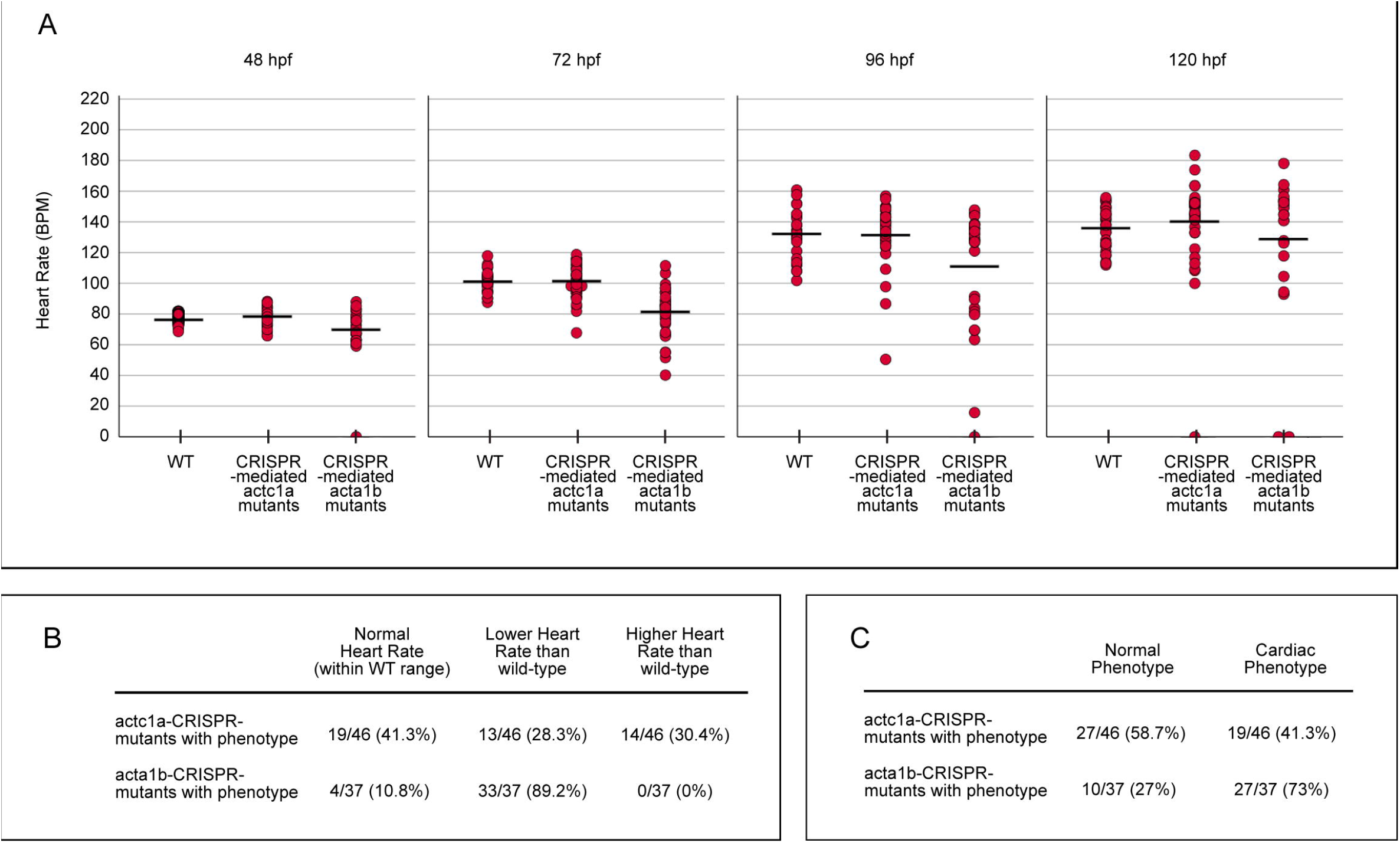
Heart Rates of CRISPR-mediated actc1a and acta1b mutants from 48-120 hpf. The heart rates of wild-type and embryos with CRISPR-generated mutations in cardiac actins *actc1a* or *acta1b* were recorded daily from 48-120 hpf (2-5 days)(A). Individual (red circles) and average (black lines) heart rates are displayed for each genotype at every age. CRISPR-mediated *actc1a* mutants had relatively similar average heart rates compared to wild-type from 48-120 hpf (A). The majority of *acta1b*-CRISPR-mediated mutants had lower average heart rates than wild-type or *actc1a*-CRISPR mutants at all stages examined (A). Subsets of the actin-CRISPR mutants started to display abnormal heart rates at 72 hpf with unique groupings for both genotypes (B). At 72 hpf, a significant number of *actc1a* and *acta1b*-CRISPR-mediated mutants developed characteristic cardiac phenotypes (pericardial edema, enlarged chambers, degenerated hearts, etc) separate from the heart rates recorded (C).

## Discussion

Zebrafish provide an excellent system for modelling human cardiac diseases and investigating the underlying mechanisms of disease progression and treatment, although a genome duplication event unique to the teleost lineage can complicate specific gene targeting studies due to genetic compensation. Zebrafish cardiac actin genes are no exception, with four identified genes (*actc1a* on chromosomes 19 and 20, *acta1b*, and *actc2*) contributing to heart development. Based on *in situ* expression (Fig 1) and previous data [16], *actc1a* and *acta1b* are transcribed in the heart throughout development. However, *actc1a* exists as two identical genes on chromosomes 19 and 20. Sequencing of these regions reveals extreme sequence identity between the two copies of *actc1a*, making it very challenging to modify and analyse one *actc1a* isoform without having to first knockout the other.

CRISPR-mediated mutants of *actc1a* or *acta1b* displayed reduced heart rates, pericardial edema, and abnormal heart anatomy, with varying severity across individuals (Fig 3 & 4). These cardiac phenotypes suggest that both *actc1a* and *acta1b* are necessary for normal heart development and function. However, CRISPR-mediated *acta1b* mutants had a higher percentage of embryos with significantly lower heart rates and cardiac phenotypes compared to the *actc1a*-CRISPR mutants. This difference in phenotypic severity between the two CRISPR mutants may be due to genetic mosaicism created by the combination of CRISPR/Cas9 cutting and error-prone cellular repair pathways (e.g., Non-Homologous End Joining) [26,27]. The mosaicism is further complicated in *actc1a*-CRISPR-mediated mutants due to the requirement of modifying both copies of *actc1a* before observing a phenotype. Although the cardiac phenotypes of our CRISPR-mediated actin mutants are similar to other zebrafish cardiac sarcomere mutants [25,28], we cannot make conclusions about the molecular pathways involved due to the various mutations present in mosaic animals that can lead to contrasting conditions within the same tissue (e.g. hypertrophic vs dilated vs restrictive cardiomyopathy [29]).

The objective of this study was to determine the best ortholog among the zfactc genes for introducing and modelling human ACTC mutations. *Acta1b* is expressed in the heart during embryogenesis, represents a single efficient gene target, and CRISPR-mediated *acta1b* mutants displayed distinct cardiac phenotypes, making it an ideal candidate for characterizing the phenotype and disease mechanism of specific human actin mutations without compensation from an identical gene copy. This work establishes a foundation for modelling human actin mutations in zebrafish, and future research will focus on introducing human cardiac actin mutations into *acta1b* and characterizing the disease, as well as exploring treatment methods.

## Conclusions

Zebrafish provide a uniquely advantageous animal model for studying mutations that cause cardiomyopathy, as they can survive embryogenesis without a functional heart. However, the duplication of the teleost genome can complicate the investigation of specific cardiac mutations, especially when dealing with nearly identical paralog sequences and genetic compensation. In this study, we have determined that the zebrafish cardiac actin gene, *acta1b*, is the optimal candidate for introducing and studying human actin mutations identified in patients with heart disease.

